# Characterization of an enigmatic tubular ultrastructure in the bacterial defensive symbiont of the Asian citrus psyllid

**DOI:** 10.1101/2025.02.24.640011

**Authors:** Chihong Song, Junnosuke Maruyama, Kazuyoshi Murata, Toshinobu Suzaki, Atsushi Nakabachi

## Abstract

“*Candidatus* Profftella armatura” (Betaproteobacteria) is a unique organelle-like defensive symbiont harbored intracellularly within the symbiotic organ of a devastating citrus pest, the Asian citrus psyllid *Diaphorina citri* (Insecta: Hemiptera). Our previous two-dimensional transmission electron microscopy identified an unprecedented ultrastructure that appeared tubular in *Profftella*, but their detailed architecture, three-dimensional arrangement, and components were unknown. To address these issues, this study conducted serial block-face scanning electron microscopy, high-voltage electron tomography, and fluorescence *in situ* hybridization. The results revealed that highly elongated (2.8–136 µm observed), string-shaped *Profftella* cells contain tubes of various numbers (1–43 per cell observed) and length (up to 45 μm observed), depending on the cell length. The tubes were evenly distributed throughout the cells, occupying an average of 6.3% of the total cell volume. Each tube consisted of five or six thin fibers twisted into a right-handed helix, maintaining a consistent diameter of approximately 230 nm along its entire length. Even without fixation or embedding, the tubes retained their shape under high vacuum conditions in electron microscopes, demonstrating their remarkable stability and robustness. These findings suggest that the tubes may help provide mechanical stability to the highly elongated and potentially vulnerable *Profftella* cells. Further analysis showed that the tubes are closely associated with ribosomes, suggesting a role in protein synthesis. Overall, these results offer new insights into the structural and functional evolution of bacteria, with potential implications for developing novel pest control strategies.

**Significance statement:** Bacteria generally have simple intracellular structures, lacking organelles. Here, we report a highly elongated, elaborate organelle-like structure with a tubular shape in a bacterial symbiont of an important agricultural pest. The tube, which exhibits high durability and robustness, may contribute to the mechanical stability of the notably elongated symbiont cell. They are closely associated with numerous ribosomes, suggesting a potential role in gene expression. These findings not only enhance our understanding of bacterial evolution but may also provide clues for developing novel strategies for pest control.

## Introduction

In contrast to eukaryotes, bacteria typically have simpler intracellular structures, lacking the elaborate intracellular membrane systems and organelles found in eukaryotes (1). Certain ultrastructures, occasionally referred to as ‘bacterial organelles,’ such as chromatophores (2), carboxysomes (3), metabolosomes (4), and magnetosomes (5), are found in specific bacterial lineages (1). However, in most bacteria, the easily observable intracellular structures are limited only to ribosomes and, in some cases, nucleoids (6). In this context, we recently found an unprecedented ultrastructure with a seemingly tubular shape in a bacterial symbiont of a sap-sacking insect, the Asian citrus psyllid *Diaphorina citri* (Hemiptera: Sternorrhyncha: Psylloidea: Psyllidae) (7).

*D. citri* is a notorious agricultural pest that transmits “*Candidatus* Liberibacter” spp. (Alphaproteobacteria: Rhizobiales, hereafter *Liberibacter*), the pathogens of the most destructive and incurable citrus disease, huanglongbing (8, 9). *D. citri* possesses a symbiotic organ called the bacteriome (10–12), which harbors two transovarially transmitted obligate intracellular bacterial mutualists, “*Candidatus* Carsonella ruddii” (Gammaproteobacteria: Oceanospirillales, hereafter *Carsonella*) (13–15) and “*Candidatus* Profftella armatura” (Betaproteobacteria: Burkholderiales, hereafter *Profftella*) (14, 16–18). *Carsonella,* the primary symbiont conserved in Psylloidea (13–26), is a typical nutritional symbiont providing the host with essential amino acids (14, 15, 17, 24, 26, 27) scarce in the phloem sap diet (28, 29). On the other hand, *Profftella*, a secondary symbiont found exclusively in *Diaphorina* spp. (14, 16, 17), is a unique, organelle-like, versatile symbiont whose primary role appears to be protecting the holobiont (host-symbiont complex) from natural enemies by synthesizing the bioactive polyketide diaphorin (14, 17, 30–33). The *Profftella* genome is drastically reduced to 460 kb, with 15% dedicated to gene clusters for synthesizing diaphorin (14). Diaphorin is present in *D. citri* at concentrations as high as 2–20 mM, depending on its developmental stage (32). It is inhibitory to eukaryotes (14, 30, 31) and various bacteria, including *Bacillus subtilis* (Firmicutes: Bacilli: Bacillales), but promotes the growth of a limited number of bacterial species, including *Escherichia coli* (Gammaproteobacteria: Enterobacterales) (33, 34). Cell-free gene expression analyses suggest that the ribosome is a target for both the inhibitory and promoting effects of diaphorin on bacteria (35, 36). This unique ability of diaphorin to modulate bacterial vital activities may influence the microbiota of *D. citri* and potentially affect the transmission of *Liberibacter* pathogens (33). Similar to other hemipteran insects (37–48), there is growing evidence that interactions among the host psyllid, bacteriome-associated obligate mutualists, facultative symbionts, and plant pathogens play critical roles in psyllid biology and host plant pathology (9, 12, 18, 49–52).

Despite its importance for understanding *Profftella*’s unique functions, evolution, and interactions with the host and other microbes, *Profftella*’s ultrastructures remain largely uncharacterized. Although our two-dimensional (2D) transmission electron microscopy (TEM) analysis showed that *Profftella* cells contain a seemingly tubular ultrastructure unprecedented in other bacteria (7), its detailed architecture, three-dimensional (3D) arrangement within the host cell, and components remained unknown. To address these questions, in the present study, we first examined the fine 3D ultrastructure of the tube using serial block-face scanning electron microscopy (SBF-SEM) and ultra-high voltage electron microscopy (HVEM) tomography. SBF-SEM is a 3D imaging technique that generates 3D images by stacking serial section images, enabling high-resolution 3D visualization of the internal structures of biological samples (53). Electron tomography, utilizing 1000 kV HVEM, enables high-resolution 3D observation of biological specimens as thick as 1 μm, owing to the high penetrating power of the electron beam (54). As the putative spiral structure of the tube was somewhat similar to that of the nucleoid previously reported in *Bdellovibrio bacteriovorus* (Bdellovibrionota: Bdellovibrionia: Bdellovibrionales) (55) and *Synechococcus elongatus* (Cyanobacteriota: Cyanophyceae: Synechococcales) (56), we further analyzed the localization of DNA and RNA by performing DNA staining along with fluorescence *in situ* hybridization (FISH) using primers targeting the 16S ribosomal RNA (rRNA) of the bacteriome associates.

## Results

### Observation of *Profftella* and its tubes by optical microscopy and 2D TEM

To obtain an overview of *Profftella*, cells of *Profftella*, which are tightly packed in the syncytium of the host bacteriome, were released from the dissected bacteriome and directly observed without fixation, using differential interference contrast (DIC) microscopy (Fig. 1*A*). The cells were large and elongated, exhibiting various lengths and apparent widths of 2–5 µm. The tubular structures found in our previous 2D TEM analysis were also detected by optical microscopy, demonstrating that each *Profftella* cell contains numerous tubes of various lengths. Subsequently, the *Profftella* cells were lysed with detergent and dried on TEM grids without any additional processing, and the internal tubes were observed using TEM. The tubes without chemical fixation and resin embedding retained their shape even after dehydration and exposure to high vacuum conditions in the electron microscope, demonstrating their exceptional structural stability and robustness (Fig. 1*B*). The surface of the tubes showed a striated pattern (Fig. 1*C*). Their tilt series images were indicative of a right-handed helical configuration (Movie S1). The TEM analysis of *Profftella* cells within the syncytium following chemical fixation, dehydration, resin embedding, and ultrathin sectioning revealed that *Profftella* cells were densely packed within the host cells, with minimal host cell organelle in the spaces between *Profftella* cells (Fig. 1*D*). The tubes had a consistent thickness of 229.34 ± 9.57 nm (n=25) and were hollow, exhibiting a low electron density (arrowheads in Fig. 1*E* and *F*). It is important to note that sample preparation processes for electron microscopy, such as dehydration and embedding, often lead to sample shrinkage (57). Thus, the *in vivo* thickness of the tube may be slightly larger than what is observed. Given the limitations of conventional 2D TEM analysis in providing a comprehensive view of *Profftella*’s overall and fine-scale morphology, 3D reconstructions were subsequently performed.

**Fig. 1.**
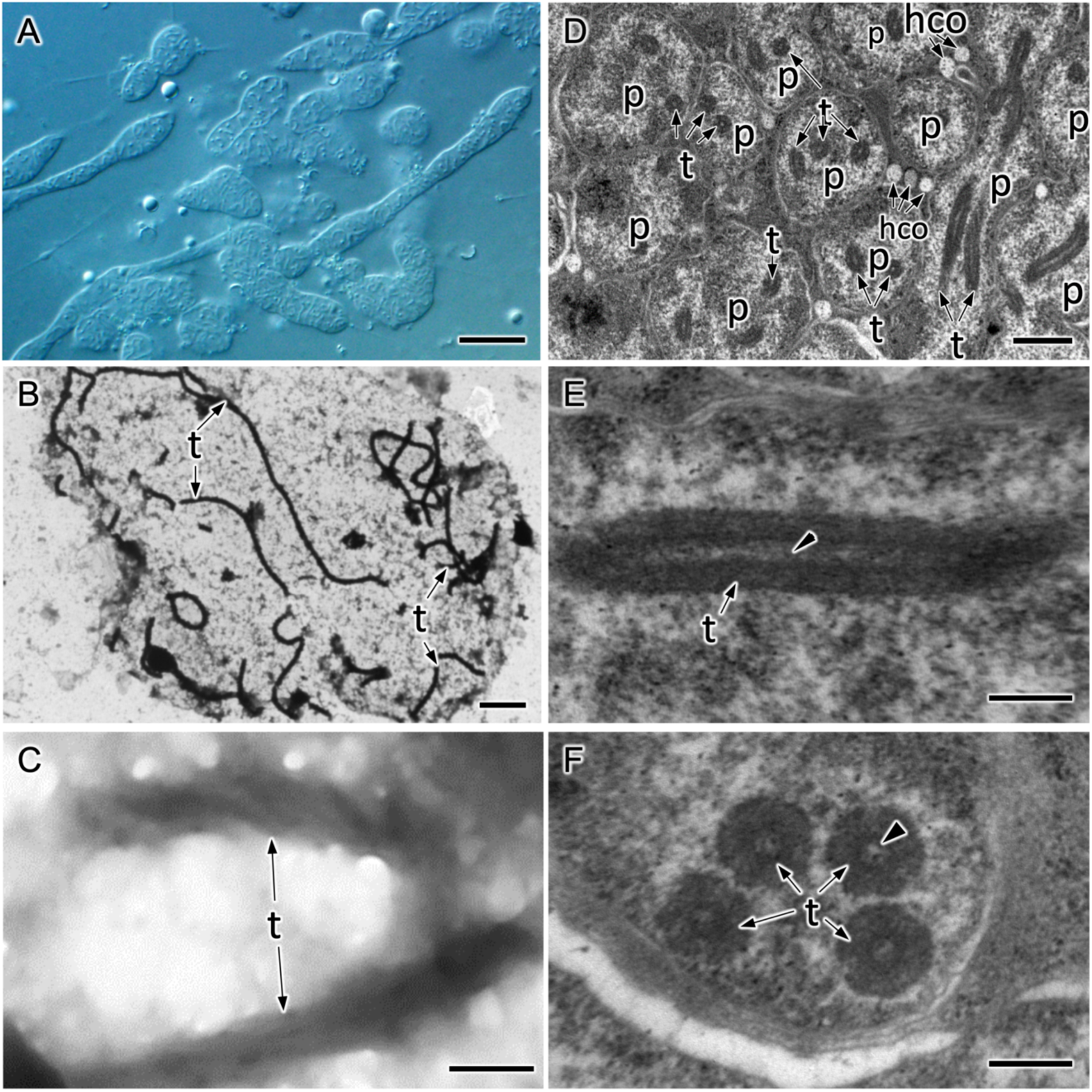
*Profftella* and tubes observed via optical microscopy and 2D TEM. (*A*) Isolated *Profftella* cells observed under an DIC microscope. (*B* and *C*) Tubes in ruptured but unfixed *Profftella* observed by TEM. (*D–F*) TEM observation of ultrathin sections of chemically fixed and resin-embedded *Profftella* cells and tubes. Numerous *Profftella* cells are packed in a syncytium of the *D. citri* bacteriome (*D*). Longitudinal view of a tube (*E*). Cross-sectional view of the tubes (*F*). p, *Profftella*; t, tube; hco, host cell organelles. Scale bars: 10 μm in (*A*), 2 μm in (*B*); 200 nm in (*C*), (*E*), and (*F*), 1 μm in (*D*).

### 3D structural analysis of Profftella and its tubes

The resin-embedded bacteriome of *D. citri* was imaged using SBF-SEM, generating approximately 200 consecutive images, each 100 nm thick. Two datasets were acquired from the same resin block. These images provided 3D information covering a volume of 20×20×20 μm³, enabling the 3D reconstruction of all *Profftella* cells and tubes within this volume (Fig. 2*A* and *E*, and Movie S2 and S3). A representative cell from each of the two consecutive image series is shown in Fig. 2*B* and *F*. These cells, measuring 49 μm and 81 μm in length, respectively, displayed an entangled appearance with no discernible pattern (Fig. 2*C* and *G*). The tubes within these cells varied in length (Fig. S1 and S2) and were uniformly distributed throughout the cytoplasm, without any specific directional preference. They occupied the intracellular space in a disordered, entangled manner and were occasionally observed in folded or coiled configurations (Fig. 2*C-D* and *G-H*, and Fig. S1 and S2). Despite the limited number of 200 consecutive images, 13 *Profftella* cells were fully reconstructed along their entire lengths (Fig. 2 and 3), ranging from 2.8 μm to 81 μm (Fig. 3*L*). The shortest cell (2.8 μm) contained only two short tubes (Fig. 3*A*), while another cell with a single tube had a volume of approximately 7.2 μm³, making it the smallest cell observed (Fig. 3*B*). Cell volume was proportional to length, with a uniform cross-sectional area of 2.1 μm², as determined through least squares regression analysis (Fig. 3*M*). Based on this, the approximate diameter of the cells was calculated to be 1.63 μm.

**Fig. 2.**
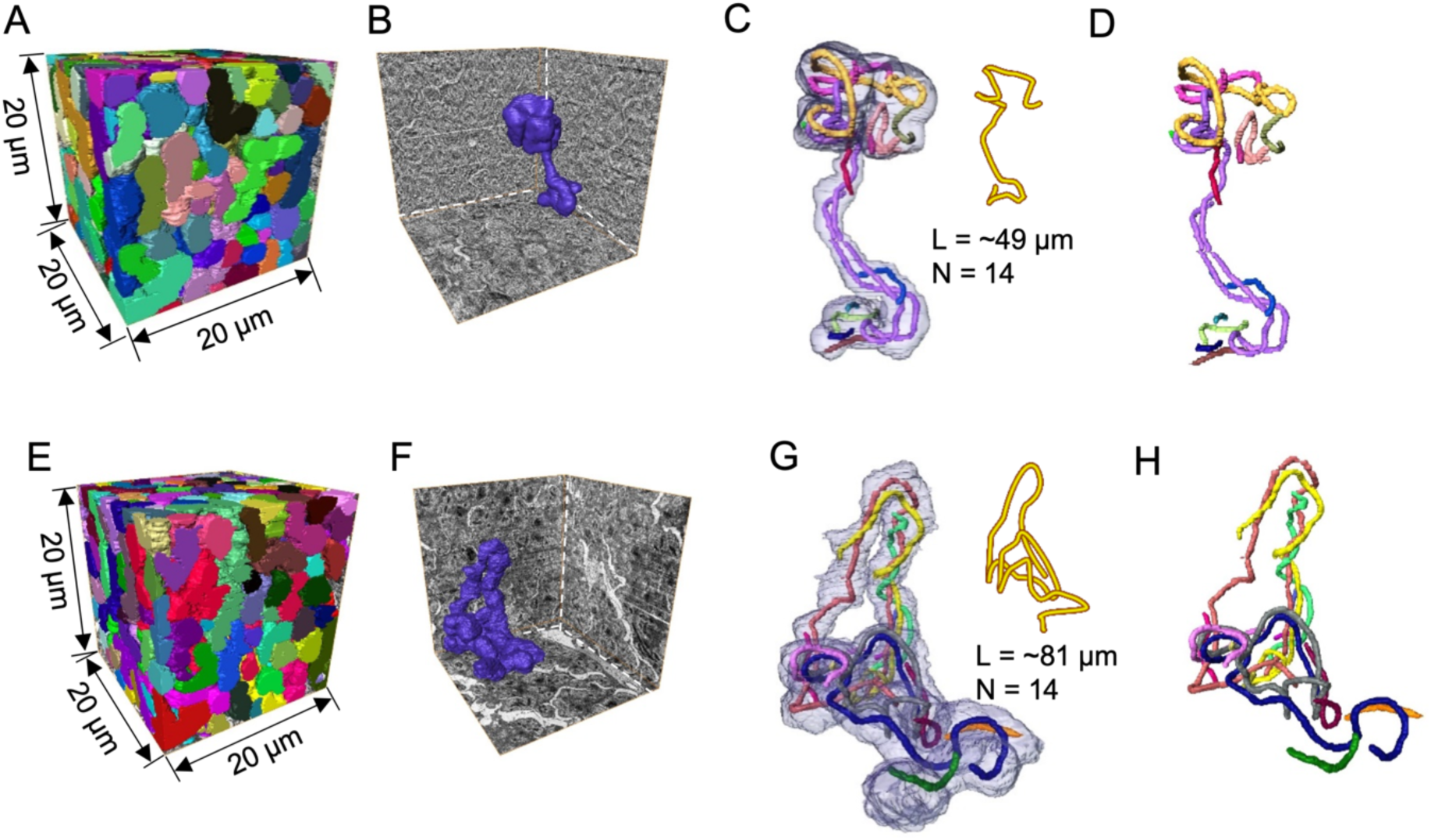
Two independent 3D reconstruction data sets including representative *Profftella* cells. (*A* and *E*) Reconstructions from 200 consecutive 100 nm-thick images obtained through SBF-SEM, showing all *Profftella* cells within each cubic region. Each side of the cubic blocks is 20 μm in length. Different *Profftella* cells are colored differently. (*B* and *F*) These images show representative *Profftella* cells where the entire cell is contained within the respective resin blocks in (*A* and *E*). The planes (XY, XZ, YZ axes) of the cubic region are also displayed. (*C* and *G*) Enlarged views of each *Profftella* cell shown in (*B* and *F*). The cell surfaces are displayed transparently, and numerous internal tube structures are color-coded for distinction. The centerline tracings of the cells, shown in the upper right corner of each figure, indicate that the cells are single-stranded. The length of each *Profftella* cell was approximately 49 μm (*C*) and 81 μm (*G*). (*D* and *H*) Multiple tube structures present inside each cell shown in (*C* and *G*). All tubes in (*D* and *H*) are individually displayed in Fig. S1 and S2.

**Fig. 3.**
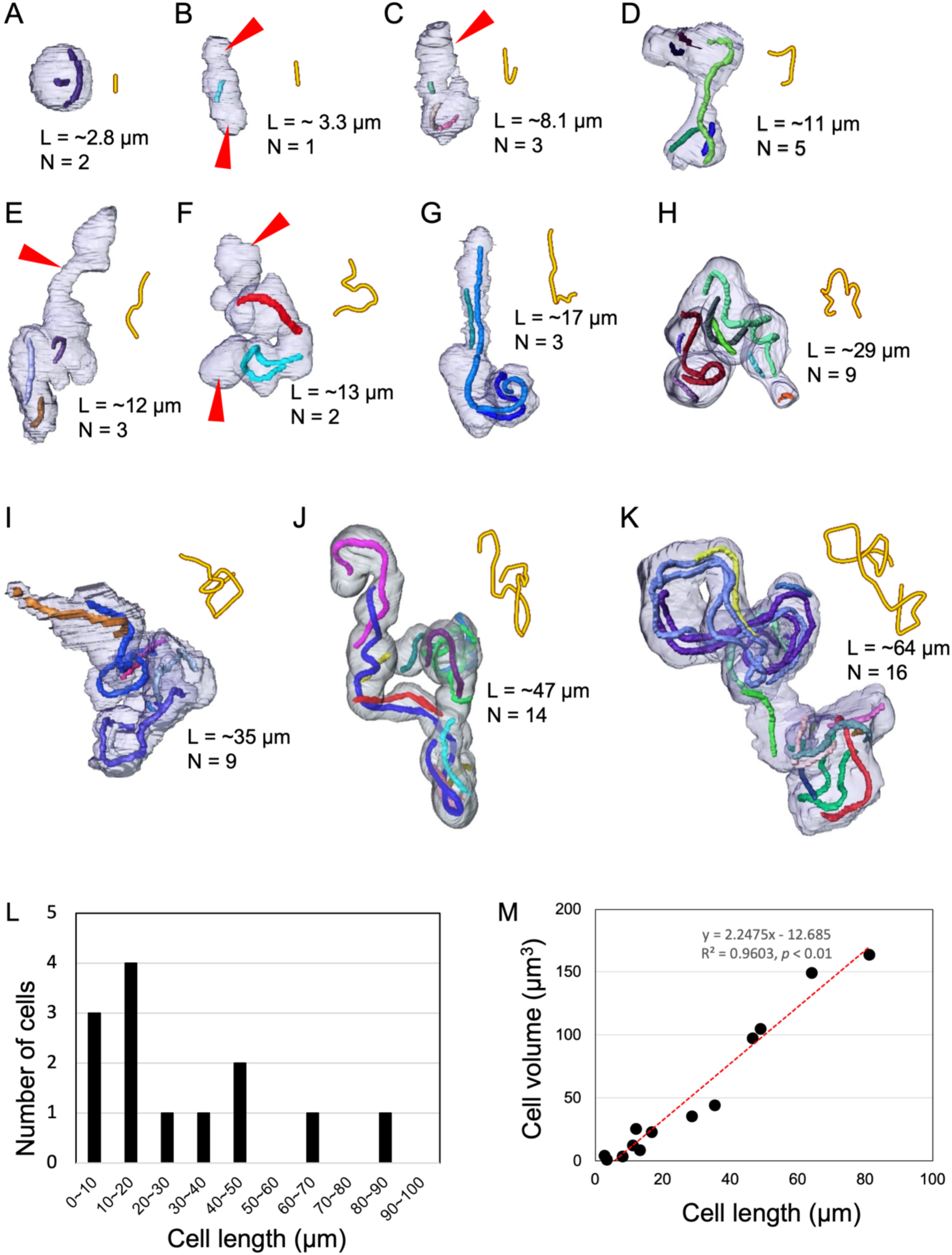
Fully-reconstructed *Profftella* cells. (*A–K*) 11 fully reconstructed cells, arranged in order of increasing *Profftella* cell length. Each figure shows the cell outline along with the tube-like structures contained (different colors represent distinct tube structures). The top-right corner of each figure displays the centerline tracing of the respective cell. Based on the centerline tracings, the cells appear as single filaments. The red arrowheads indicate regions within the cell where there are no tubes. (*L*) Length distribution of the 11 cells shown in *A–K* and the 2 cells in Fig. 2*C* and *G*. (*M*) Correlation between cell length and cell volume.

Owing to the size limitations of the analyzed volume, many cells could not be fully reconstructed. However, those exceeding 20 μm in length (based on the size of the consecutive images, see Fig. 2*A* and *E*) are shown in Fig. 4. The fully reconstructed cells were relatively small and confined to the limited imaging volume, allowing for complete visualization. In contrast, cells that could not be fully reconstructed were either larger or more widely distributed, preventing complete capture within the restricted volume (Fig. 4). The cell shown in Fig. 4*E* had a zigzag shape with four bends and a length exceeding 100 μm. The longest cell observed had a reconstructed length of 136 μm, indicating that its actual length is likely even longer (Fig. 4*F*). This cell uniquely displayed a single branching point, suggesting that larger cells may branch or divide into two segments (Fig. 4*F* and *G*). It also contained the highest number of tubes (43 in total) among all observed cells (Fig. 4F). At the branching point, a single long tube extended across both branches (Fig. 4*H*), confirming that the segmentation and rendering process did not erroneously identify two separate cells as one.

**Fig. 4.**
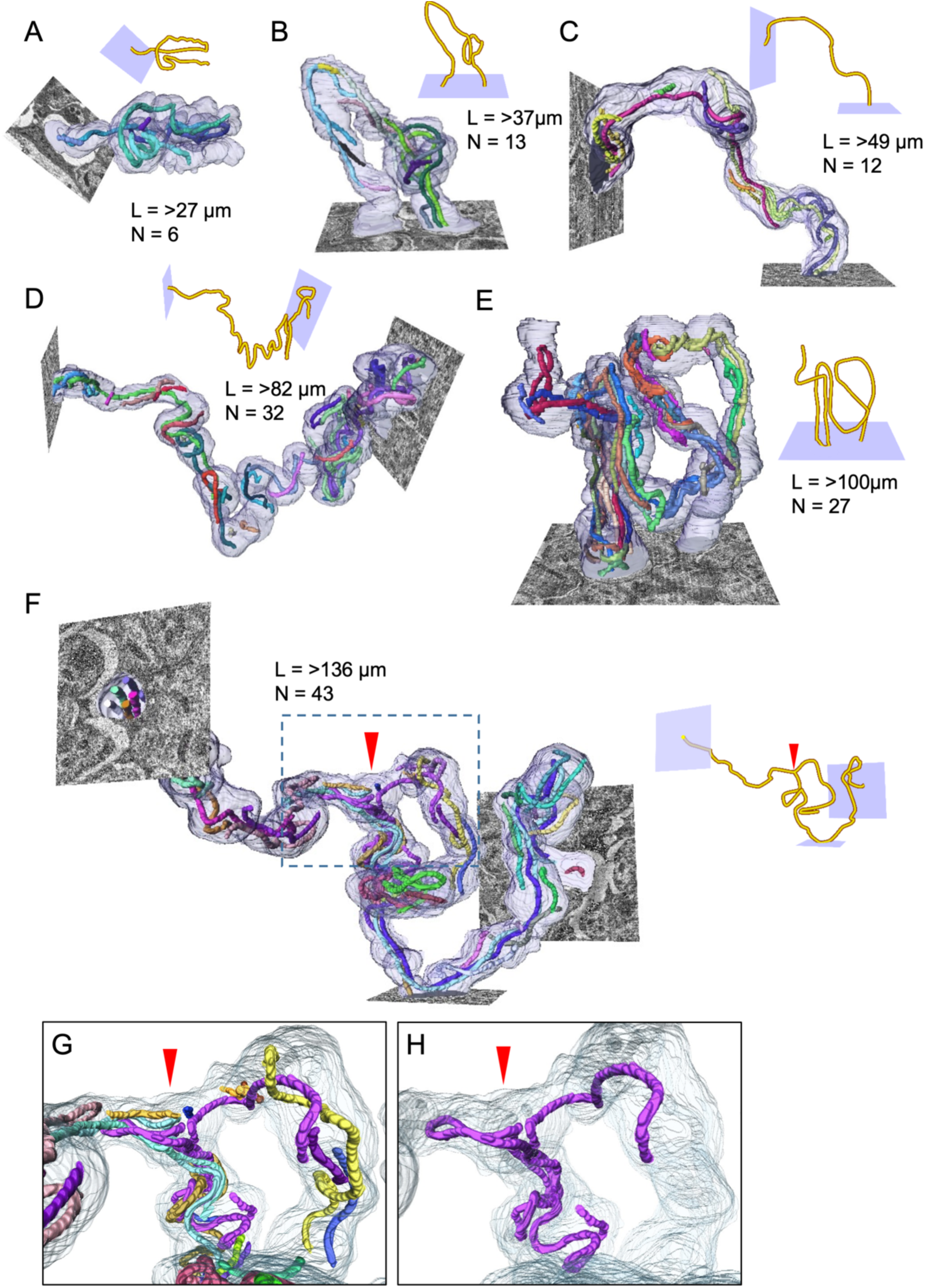
Reconstruction of *Profftella* cells that were not fully contained within the analyzed cubes. (*A–F*) Due to the data size limitations, some of the entire cells could not be fully reconstructed, but the six incomplete cells longer than 20 μm are shown. The cells are ordered by increasing length. Cross-sectional images of the block showing the ends of the *Profftella* cells are also included. For each cell, the centerline tracing, the cell length (L, not the full length), and the number of tubes (*N*) are indicated with the corresponding reconstructed images. The arrowhead indicates the branching point of the cell in (*F*). (*G*) Magnified view of the branching point of (*F*). (*H*) One of the tubes at the branching point extends on both branches.

All reconstructed cells contained a total of 230 tubes, with considerable variability in their lengths. Due to the 100 nm section thickness, it was impossible to measure tubes smaller than this precisely. The longest tube observed in the cell shown in Fig. 4*F* was approximately 45 μm in length (Fig. 5*A*). Notably, as cell volume increases to approximately 80–90 μm^3^, the internal tube occupancy increases proportionally, after which it stabilizes at around 7% of the cell volume (Fig. 5*B*). This is consistent with our observation of the reconstructed cell structures shown in Fig. 3*A-K*. In smaller cells, tube-free regions were observed at the ends (red arrowheads in Figs. 3*B–C* and *E–F*), indicating a lower tube volume fraction (Figs. 3*A–G*). In contrast, larger cells contained densely packed tubes distributing throughout the cytoplasm, indicating a consistent volume fraction (Figs. 3*J–K* and 4*C–F*). This suggests that total volume of the tube is determined by the size of the cell (Fig. 5*B*). Additionally, no cells completely lacking tubes were observed, implying that tubes arise from the division of pre-existing tubes and grow along with the cells as they enlarge.

**Fig. 5.**
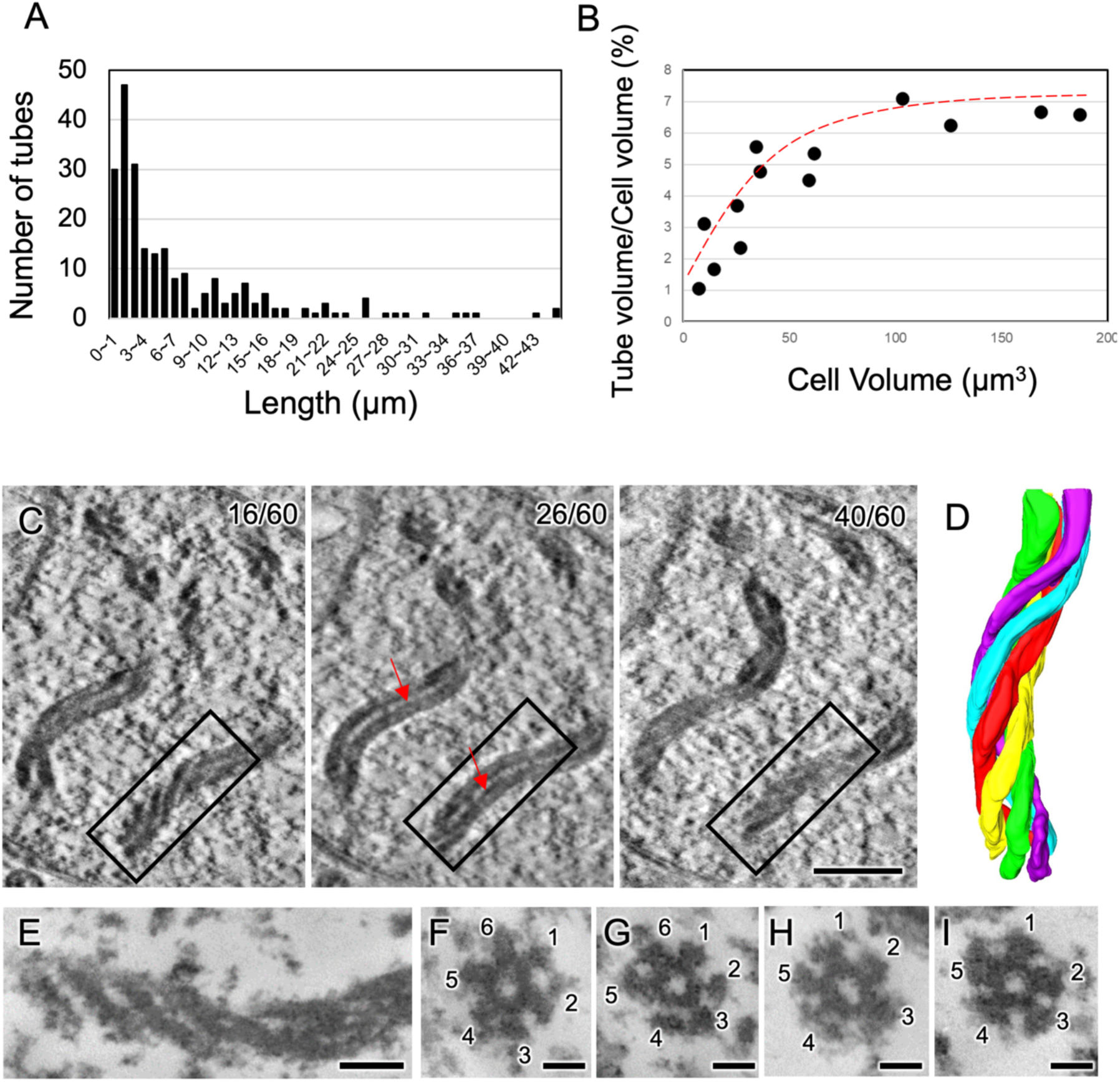
Length, volume, and nanostructure of the tubes in the cell. (*A*) Distribution of the tube length (N=230). (*B*) Relationship between cell volume and total tube volume (N=13). (*C*) Tomographic slices of the 8 nm thick Z-plane reconstructed by HVEM, showing the ultrastructure of the tubes within *Profftella* cells (see also Movie S4). The left, middle, and right panels represent the far slice of the tubes (16th of 60 slices), the center slice of the tubes along with internal hollows (indicated by red arrows, 26th of 60 slices), and the near slice of the tubes (40th of 60 slices), respectively. (*D*) A volume-rendering image of the tube corresponding to the rectangular region in (*C*) (see also Movie S5), showing five fibers twisted in a right-handed helical configuration. (*E–I*) TEM images of the tubes observed in 100 nm thick sections after detergent treatment, showing a longitudinal section of the tube displaying obliquely arranged sub-filaments (*E*), and cross-sections of the tubes (*F-I*) indicating that the outer wall of the tube is composed of five or six sub-filaments. Scale bars are 500 nm in (*C*) and 100 nm in (*F-I*).

To analyze the finer structure of the tubes, relatively thick sections (~500 nm) were prepared for electron tomography using HVEM. Tomography data revealed that five fibers were twisted together in a right-handed helical configuration to form the tube (Fig. 5*C* and *D*, and Movie S4–S6). This helical structure was also observed in unembedded bare tubes as described above (Movie S1). The central region appeared hollow with no internal filling. To more accurately determine the number of fibers, specimens were treated with Triton X-100, fixed, dehydrated, resin-embedded, and sectioned for TEM observation (Fig. 5*E–I*). Cross-sectional analysis revealed that the tubes were composed of five or six fibers (Fig. 5*F-I*). Out of ten clearly observed cross-sections of the tubes, eight contained six fibers and two contained five fibers, suggesting that the six-fiber type is more common, while the five-fiber type is less frequent.

### Tubes contain ribosomes

FISH analysis using probes specific for the 16S rRNA of *Profftella* or *Carsonella*, along with DNA staining using Hoechst 33342, clearly distinguished between *Profftella* and *Carsonella* released from the *D. citri* bacteriomes (Fig. S3). DIC microscopy demonstrated that tubes are present only in *Profftella,* not in *Carsonella*, which is consistent with our previous TEM observations (7). The probe specific for *Profftella* 16S rRNA colocalized with the tubes in *Profftella* (Fig. 6*A* and Movie S7), while DNA signals were distributed widely within the *Profftella* cells and did not align with the tubes. These results suggest that ribosomes, but not DNA, colocalize with the tubes. To assess the possibility that the probe also binds to the genomic DNA (16S rRNA gene) of *Profftella*, nuclease treatment was further performed before hybridization (Fig. 6*B* and Movie S8 and S9). DNase treatment had no effect on the colocalization of the probe with the tubes (Fig. 6*B*, Movie S7), whereas RNase treatment abolished the probe signal (Fig 6*C* and Movie S8). This supports the hypothesis that ribosomes, not DNA, colocalize with the tubes, as RNase treatment is expected to degrade ribosomes by digesting rRNA, which accounts for approximately 65% of the ribosome’s molecular weight (58). These results indicate that the tube is not a nucleoid but rather a structure that contains ribosomes as its components. To further investigate the structural relationship between the tubes and ribosomes, we performed TEM observations after RNase treatment. In the control group, granular structures attached to the tubes were observed (Fig. 6*D*), whereas in the RNase-treated group, these granular structures were absent, although the morphology of the tubes remained almost unchanged (Fig. 6*E*). These findings suggest that ribosomes are present within the tube but are not essential for its structural integrity.

**Fig. 6.**
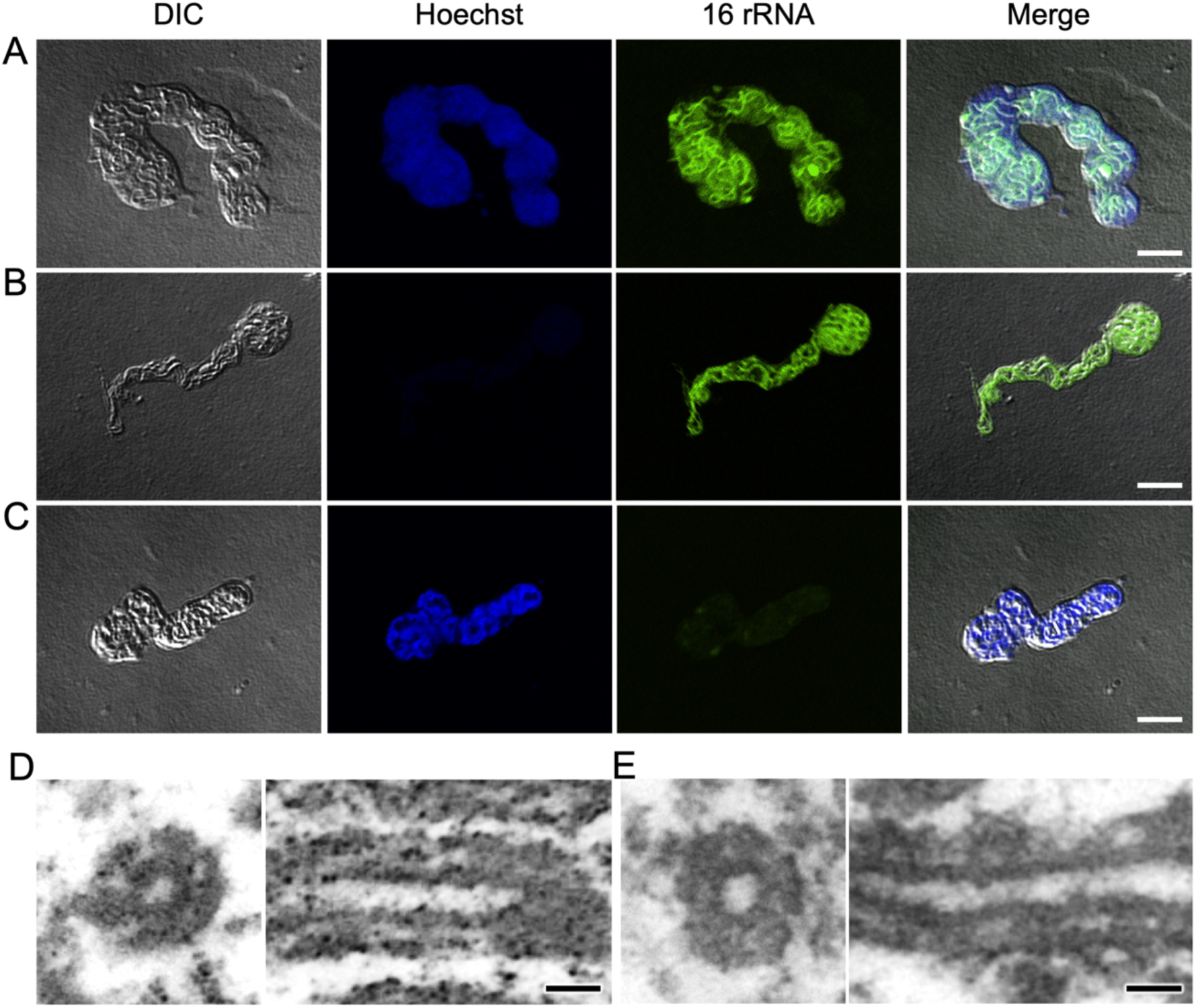
FISH and TEM observations of the tubes with and without nuclease treatments. (*A-D*) FISH images of *Profftella* cells: *Profftella* cell without nuclease treatment (*A*); *Profftella* cell with DNase treatment (*B*); *Profftella* cell with RNase treatment (*C*). In addition to these Alexa 488 (16S rRNA of *Profftella*) signals, DIC, Hoechst 33342 (DNA) images are also shown along with merged images indicating 16S rRNA colocalization with the tubes. (*D*) TEM images of the tubes without RNase treatment: cross-sectional (left) and longitudinal (right) views. (*E*) TEM images of the tubes with RNase treatment: cross-sectional (left) and longitudinal (right) views. Scale bars are 5 μm in (*A-C*) and 100 nm in (*D* and *E*).

## Discussion

This study revealed that *Profftella* cells contain tubes of various lengths (up to 45 μm) and numbers (ranging from 1 to 43 per cell). Each tube consisted of five or six thin fibers twisted into a right-handed helix, with a consistent diameter of approximately 230 nm along its entire length. This twisted, multi-fiber structure likely enhances the tubes’ durability and flexibility compared to a single-fiber design. It closely resembles the construction of steel wire ropes or the thick and sturdy ropes used in “tug-of-war,” where multiple thinner strands are twisted together to enhance strength, durability, flexibility, and resilience (59). The tubes with this multi-fiber structure, even without fixation or embedding, retained their shape after being treated with detergent, dried, and placed under the high vacuum conditions of electron microscopes, demonstrating their remarkable stability and robustness (Fig. 1*C* and Movie S1). These findings suggest that the tubes, like the cytoskeletons of eukaryotes (60), may help provide mechanical stability to the highly elongated and potentially vulnerable *Profftella* cell while maintaining its flexibility. Additionally, the intracellular occupancy of the tubes increased proportionally with the cell volume up to 80–90 μm^3^ and then stabilized at approximately 7% (Fig. 5*B*). This indicates that tube volume is regulated in relation to cell length, implying that *Profftella* optimizes the tube volume to maintain structural stability across various sizes and shapes. Cytoskeletons not only provide physical support for the cell structure but also facilitate the efficient and targeted transport of various substances, including nutrients and wastes, in large cells (61, 62). Thus, the tubes in *Profftella* may also similarly serve as scaffolds for substance transport in these highly elongated *Profftella* cells.

This study also demonstrated that *Profftella* cells are highly elongated, exhibiting considerable variability in cell length, ranging from 2.8 μm to over 136 μm. This finding is consistent with our previous observation that *Profftella* cells are spherical when transferred from the bacteriome to the ovary but begin to elongate upon entering the oocytes, implying that *Profftella* is string-like within the bacteriome but transforms to a spherical shape before exiting into the hemocoel (18). However, as the *Profftella* cells are tightly packed in the bacteriome, their precise shape and 3D arrangement remained unclear. This study resolves that uncertainty, demonstrating that the *Profftella* cells are highly elongated within the host bacteriome. While these morphological features are atypical from a bacteriological perspective, they are not necessarily unique to insect intracellular mutualists. For instance, *Carsonella*, the primary symbiont of psyllids (15, 47, 63, 64), and “*Candidatus* Sulcia muelleri” (Flavobacteriia: Flavobacteriales, hereafter *Sulcia*), an ancient obligate mutualist of various auchenorrhynchan insects (65, 66), also exhibit notable elongation in the bacteriome. Moreover, *Carsonella* forms spherical cells before transferring from the bacteriome to the ovary, where they enter the oocyte, forming a “symbiont ball,” which is a mosaic mass of *Carsonella* and *Profftella* (18). The highly elongated shapes of *Profftella*, *Carsonella*, and *Sulcia* may reflect a survival strategy that prioritizes resource allocation toward symbiont functions, such as syntheses of nutrition and bioactive substances, rather than excessive investment in cell division. None of these symbionts retains the *ftsZ* gene (14, 15, 26, 67, 68), a highly conserved gene playing a central role in bacterial cell division (69, 70), which may partly explain the elongated morphology of these symbionts. However, the intracellular tubular structures are found only in *Profftella*. The reason for this, as well as the specific *Profftella* genes responsible for the tube formation, are yet to be elucidated.

FISH analysis using a probe targeting the 16S rRNA of *Profftella*, combined with DNase/RNase treatment and DNA staining with Hoechst 33342, demonstrated that ribosomes, but not DNA, colocalize with the tubes. This rules out the possibility that the tubes are nucleoids, as seen in *B. bacteriovorus* (55) and *S. elongatus* (56). In *B. bacteriovorus*, approximately 13% of cells exhibited twisted, multi-rod-shaped nucleoid bodies arranged in a spiral formation, with length of up to approximately 1.5 μm. These nucleoids were loosely wound spirals with diameters ranging from 50 to 200 nm along their length (55). In *S. elongatus*, the seemingly single, wavy, rod-shaped nucleoids, approximately 3 μm in length and observed only before cell division (55), varied in diameter from 100 to 500 nm (56). In contrast, the tubes in *Profftella* maintain a constant diameter of 230 nm along their entire length, and their structure is clearly visible under DIC microscopy, further supporting the idea that they are not nucleoids. In *Profftella*, the DNA signal was broadly distributed throughout the cell (Fig. 6*B*). The relatively strong DNA signal (Fig. S3*B*) in *Profftella*, compared to *Carsonella*, suggests that *Profftella* harbors numerous copies of its highly reduced 460 kb genome, as confirmed in *Carsonella*, which contains thousands to tens of thousands of highly reduced (160–175 kb) genomic copies per cell (63, 71, 72). TEM analysis following RNase treatment further supported the presence of ribosomes in the tubes, suggesting their potential involvement in gene expression. However, the analysis also indicated that ribosomes are not essential the structural integrity of the tubes, highlighting the need for further investigation to identify their primary constituents. Additional studies are required to fully elucidate the components and functions of the tubes.

## Materials and Methods

### Insects

An established colony of *D. citri*, initially collected on Amami-Oshima island, Kagoshima, Japan, was maintained on the saplings of the orange jasmine, *Murraya paniculata* (Rutaceae). The plants covered with insect-rearing sleeves were kept in incubators at 28°C with a 16:8 (light: dark) h photoperiod. Adult insects were collected from the plants using an insect aspirator and then caged in a plastic dish on ice for 5 min to immobilize them. They were sexed under a stereomicroscope, and bacteriomes were dissected from the abdominal hemocoel of adult females in phosphate-buffered saline (PBS: 0.8% NaCl, 0.02% KCl, 0.115% Na_2_HPO_4_, 0.02% KH_2_PO_4_).

### Optical microscopy of unfixed *Profftella* cells

After dissection, bacteriomes were gently crushed with forceps and repeated pipetting in PBS, and *Profftella* cells released from the syncytium of the bacteriome were placed on glass slides, covered with coverslips, and observed using DIC microscopy (BX-53; Olympus, Tokyo, Japan).

### TEM observation of the tubes without fixation

*Profftella* cells treated with detergent were observed by conventional TEM. First, bacteriomes removed from insect bodies were subjected to cell disruption by vigorous agitation in a detergent medium (0.5% Triton X-100, 5 mM MgSO_4_, and 100 mM HEPES-KOH pH 7.0) for 10 min at room temperature. Subsequently, some of the disrupted cells were placed directly onto TEM grids without fixation or any other additional processing to observe the tubes. To capture the external morphology from multiple angles, tilt series images were recorded from −60° to +60° using a JEM-2100F (JEOL, Tokyo, Japan) at an accelerating voltage of 200 kV, with the results provided as Movie S1.

### TEM observation of *Profftella*

Equal volumes of glutaraldehyde fixative (6% glutaraldehyde, 40 μM MgSO_4_, 4 mM sucrose, and 100 mM cacodylate buffer pH 7.0) were added to the sample suspensions and chemically fixed on ice for 4 h. The cells were then postfixed in OsO_4_ fixation solution (1% OsO_4_, 20 µM MgSO_4_, 2 mM sucrose, and 50 mM cacodylate buffer pH 7.0) for 1 h at room temperature. The samples were then dehydrated in an ethanol series, embedded in Spurr’s resin, and 100 nm sections were observed on a Hitachi H-7100 transmission electron microscope operating at 75 kV.

### Sample preparation for 3D electron microscopy

The specimens were initially treated at room temperature for 10 minutes with 3% glutaraldehyde in 50 mM Na-cacodylate buffer (pH 7.0), supplemented with 20 μM MgSO4 and 2 μM sucrose. Following the chemical fixation, the materials washed three times in the aforementioned buffer and were subsequently postfixed at room temperature for 30 minutes with 1% OsO4 in the same buffer. Fixed cells were then subjected to dehydration through a graded ethanol series (50%, 70%, 90%, 95%, 99%, and 100%) and embedded in Spurr’s resin at 70°C for 8 hours. Serial ultrathin sections (approximately 100 nm thick) were prepared using a diamond knife (EM UC7 ultramicrotome; Leica, Austria). The resulting ultrathin sections were stained with 3% uranyl acetate and lead citrate before examination utilizing an H-7100 transmission electron microscope (Hitachi, Japan) operating at 75 kV.

### SBF-SEM

The resin block containing the specimens was trimmed and affixed to an aluminum rivet using conductive epoxy resin (SPI Conductive Silver Epoxy; SPI Supplies and Structure Probe, Inc., West Chester, PA, USA). Subsequently, the block was coated with gold using an ion coater. SBF-SEM (ΣIGMA/VP, Carl Zeiss Microscopy, Jena, Germany; 3View; Gatan Inc., Pleasanton, CA, USA) with a back-scattered electron detector and a diamond-knife for serial sectioning was employed for slicing and imaging the specimen. The specimen was introduced into the SBF-SEM chamber, and the block face was aligned parallel to the knife edge and brought close to the knife’s height. To mitigate charging effects, the electron microscope operated at a low accelerating voltage of 1.5 kV. Serial image series were acquired in an automated manner using Gatan Digital Micrograph software, with all images captured at a size of 8192 × 8192 pixels (pixel size = 3 nm). For SEM image acquisition, a 100-nm-thick layer was automatically removed from the block face by the knife to expose a fresh surface for imaging, and this cycle of image acquisition and block face removal was repeated. Two datasets, each consisting of 200 serial images, were acquired and aligned using the IMOD software package (73). Subsequent segmentation of regions of interest was carried out using Amira version 5.4.5 (FEI Visualization Science Group, Burlington, MA, USA). The lengths of the reconstructed cells and tubes were manually measured using Amira, and their volumes were automatically calculated in Amira. A linear regression analysis using the least squares method was performed to evaluate the relationship between cell length and cell volume (Fig. 3*M*).

### High-voltage electron tomography

A 500 nm-thick section collected on a single-slot grid was subjected to imaging without staining. Data acquisition was conducted using a 1,000 kV electron microscope (H1250M, Hitachi). Tilt series were captured on a 2k×2k CCD camera (FC400, Direct Electron LP) at 2° increments within a tilt range from −60° to +60°. Following image alignment, the 3D reconstruction was performed through weighted back-projection using IMOD software (73). Segmentation was carried out using Amira version 5.4.5 (FEI Visualization Science Group, Burlington, MA, USA).

### FISH

Bacteriomes were dissected from *D. citri*, gently crushed with forceps and repetitive pipetting to release *Profftella* cells, and fixed with 4% paraformaldehyde/PBS for 90 min. After washing with PBS twice, the samples were suspended with Milli-Q water, applied to glass slides, and dried on a hot plate at 40°C. Subsequently, the hybridization buffer (20 mM Tris-HCl [pH 8.0], 0.9M NaCl, 0.01% SDS, and 30% formamide) without probes was added and pre-incubated at room temperature for 30 minutes. Samples were then incubated at room temperature overnight with hybridization buffer containing 100 nM each of the probes SSDC_127247 (5’-GACCCTCTGTATGCACCATT-3’), 5’-labeled with Alexa Fluor 488 to specifically detect 16S rRNA of *Profftella*, and Car1 (5’-CGCGACATAGCTGGATCAAG-3’), 5’-labeled with Alexa Fluor 568 to specifically detect 16S rRNA of *Carsonella*. After washing three times with PBSTx (0.8% NaCl, 0.02% KCl, 0.115% Na2HPO4, 0.02% KH2PO4, and 0.3% Triton X-100), the samples were mixed with NucBlue Live ReadyProbes Reagent (Thermo Fisher Scientific, Waltham, MA, USA), a stabilized solution of Hoechst 33342 (2’-[4-ethoxyphenyl]-5-[4-methyl-1-piperazinyl]-2,5’-bi-1H-benzimidazole), a fluorescent stain for DNA. Samples were then mounted in ProLong Gold antifade reagent (Thermo Fisher Scientific) using a coverslip and were examined using a Nikon A1 laser scanning confocal microscope. Acquired images were analyzed using NIS-elements AR Analysis 4.10 software (Nikon).

### FISH with nuclease treatment

After fixation and washing as above, the samples were treated with DNase or RNase. DNase treatment was performed at 37 °C for 30 min using 10× DNase I Buffer, 10U of Recombinant DNase I (RNase-free, Takara, Kusatsu, Japan), and 20U of RNase Inhibitor (Thermo Fisher Scientific), followed by incubation with 0.5M EDTA at 80 °C for 2 min. RNase treatment was performed at room temperature for ten minutes with 2.5 μg/μL of RNase A (Takara). After nuclease treatment, the samples were washed with PBS, suspended with Milli-Q water, applied to glass slides, dried, and FISH was performed as described above.

### TEM observation with RNase treatment

After dissection in HEPES buffer (100 mM HEPES-KOH [pH 7.0], 5 mM MgSO_4_, and 0.5% Triton X-100), bacteriomes were crushed to release *Profftella*, fixed by adding equal volumes of glutaraldehyde fixative (6% glutaraldehyde, 100 mM HEPES-KOH [pH 7.0], 4 mM sucrose, and 40 μM MgSO_4_). Subsequent sample preparation and observation by TEM were performed using the protocol described in the “TEM Observation of *Profftella*” section.

## Acknowledgments

We thank Ms. Sachiko Yamada for her valuable assistance in segmenting the SBF-SEM data. This work was supported by the Japan Society for the Promotion of Science (https://www.jsps.go.jp) KAKENHI (grant numbers 21687020, 26292174 and 20H02998 to AN), the Collaborative Study by High Voltage Electron Microscopy Program (2015-502, 2016-502) of National Institute for Physiological Sciences to AN, and a National Research Foundation of Korea (NRF) grant RS-2024-00440289 to CS. The funders had no role in the study design, data collection and analysis, decision to publish, or manuscript preparation.

## Author contributions

C.S., J.M., K.M., T.S. and A.N. performed research; and C.S., T.S. K.M. and A.N. wrote the paper.

---

**Fig. S1.**
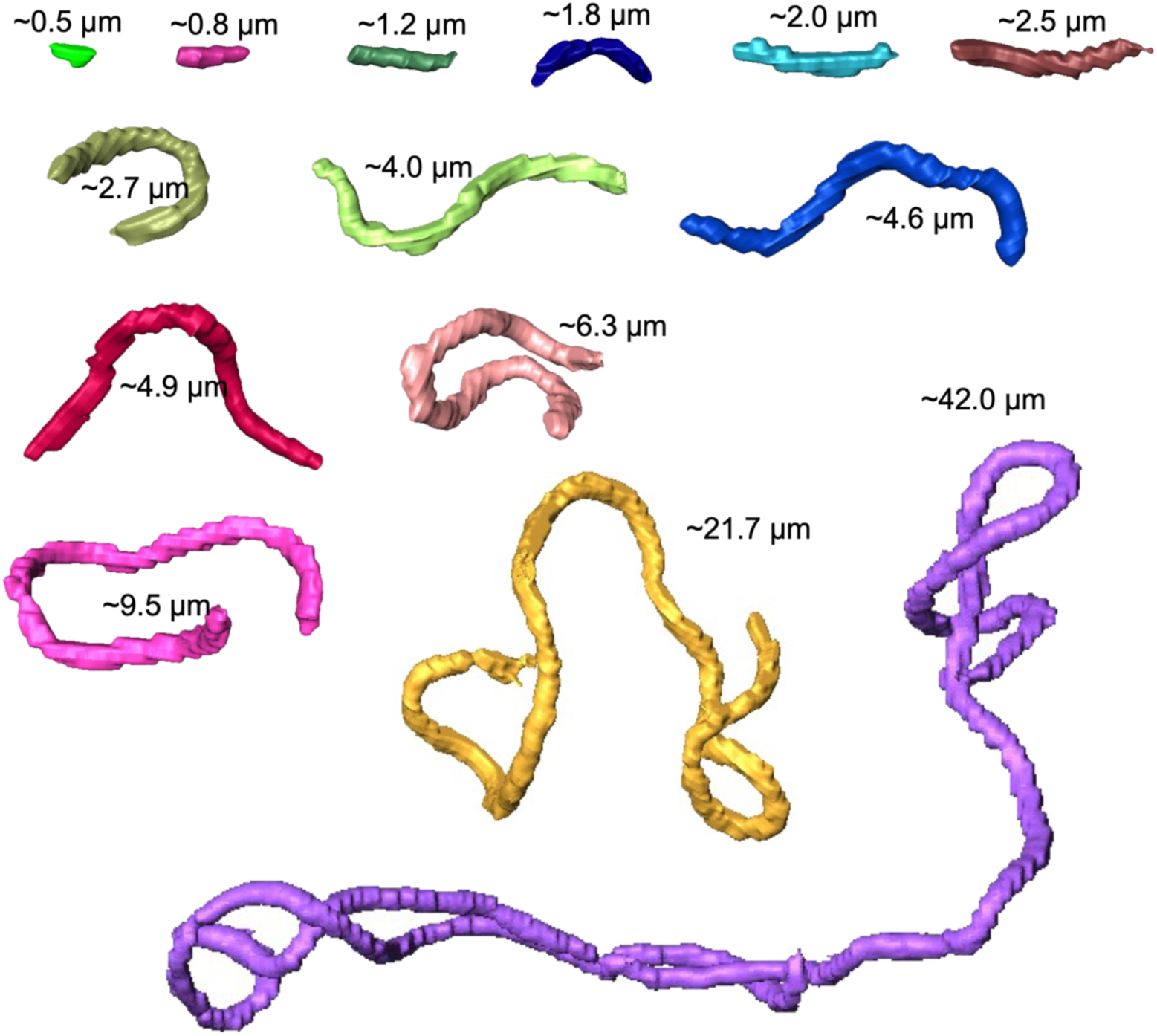
Each appearance of the tubes contained within Figure 2D.

**Fig. S2.**
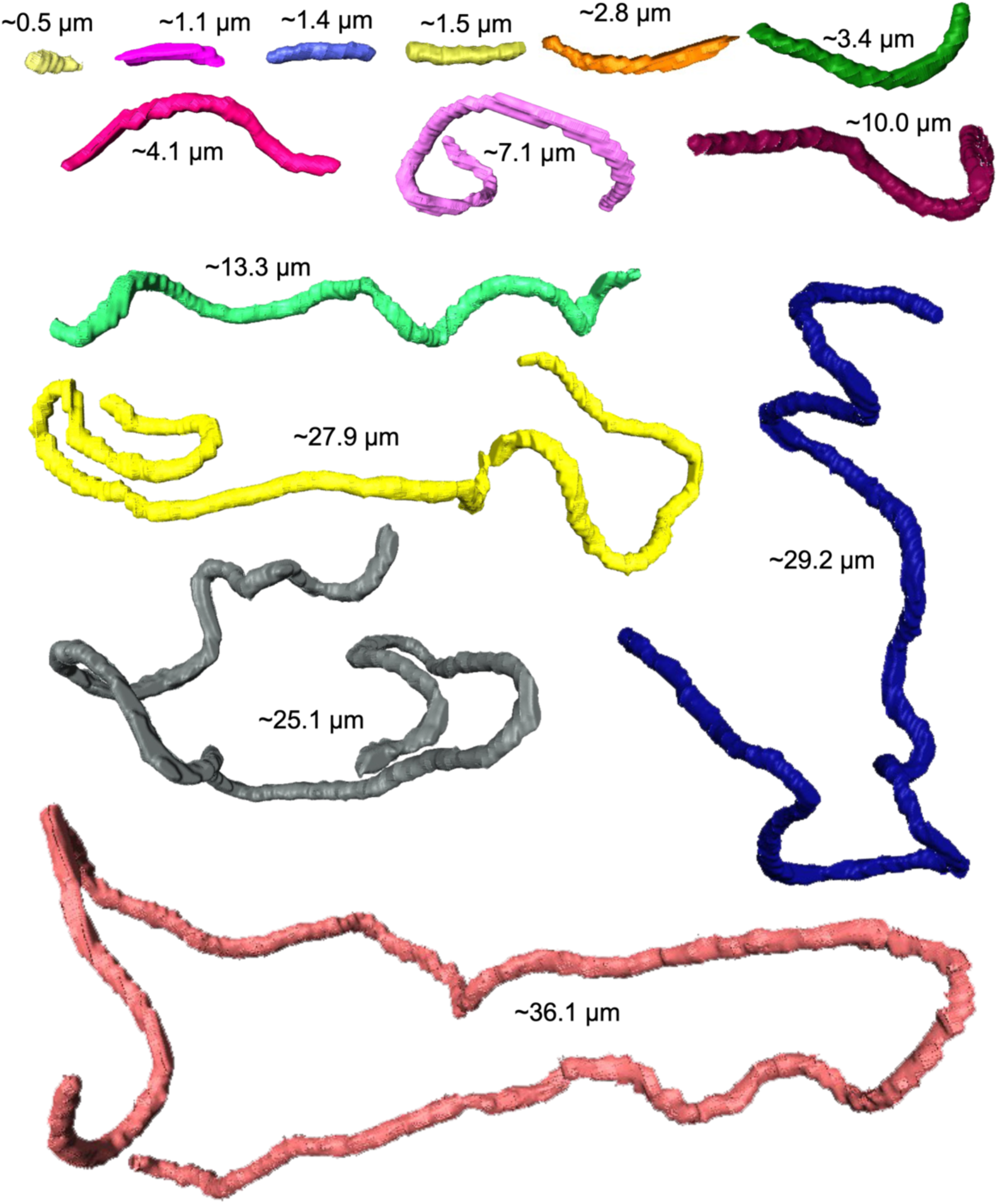
Each appearance of the tubes contained within Figure 2H.

**Fig. S3.**
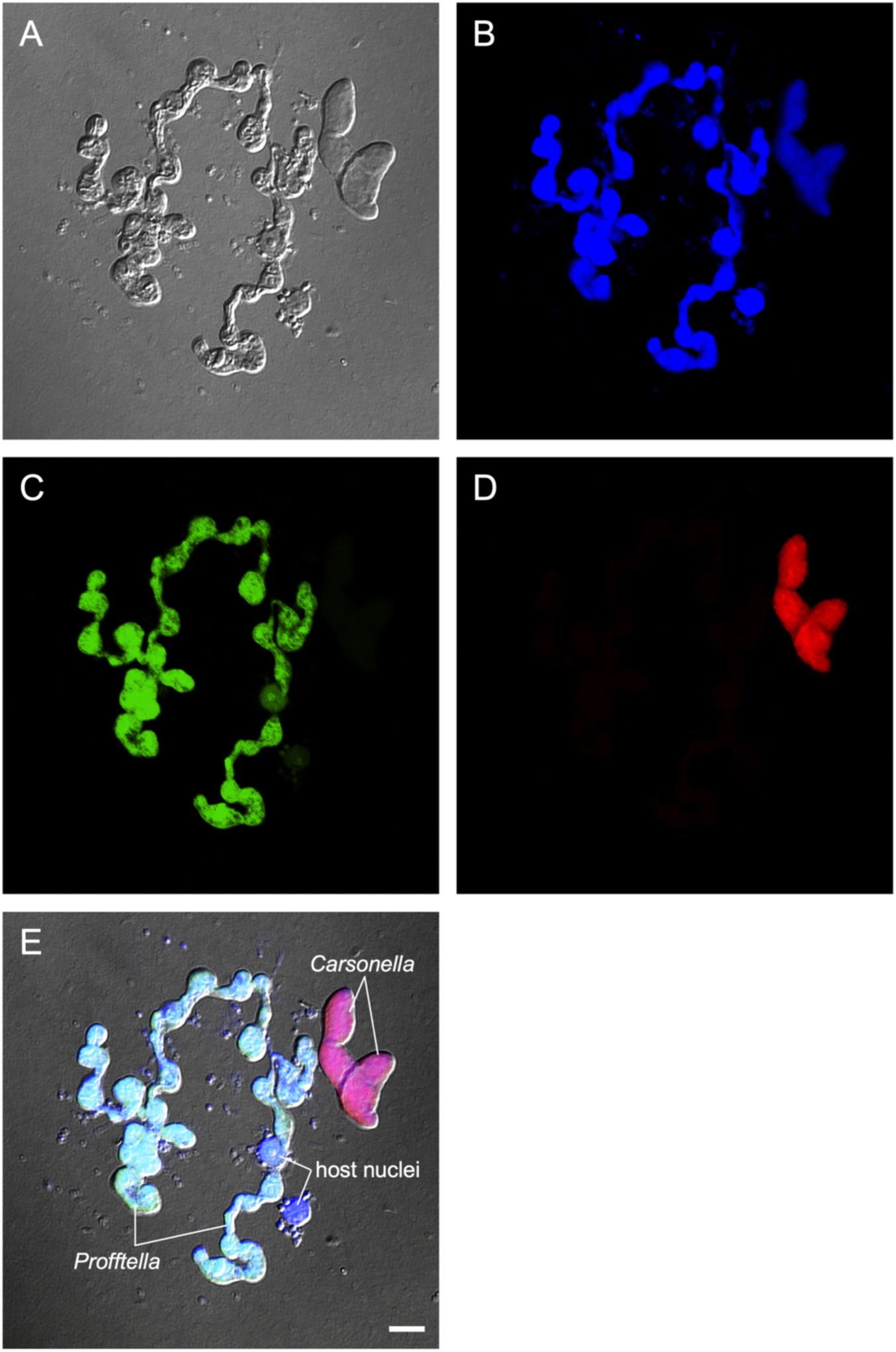
Images of FISH for Profftella and Carsonella, and the host nuclei derived from the syncytium of the *D. citri* bacteriome. (*A*) DIC image. (*B*) Signals of Hoechst 33342 showing distribution of DNA. (*C*) Signals of Alexa 488 showing distribution of 16S rRNA of Profftella. (*D*) Signals of Alexa 568 showing distribution of 16S rRNA of Carsonella. (*E*) Merged image of (*A–D*). Scale bar: 5 μm.

**Movie S1 (separate file).**

Tilt image series of detergent-treated Profftella cells observed by TEM.

**Movie S2 (separate file).**

3D reconstruction set 1 of *Profftella* cells generated by SBF-SEM.

**Movie S3 (separate file).**

3D reconstruction set 2 of *Profftella* cells generated by SBF-SEM.

**Movie S4 (separate file).**

Tilt image series of tubes observed by HVEM.

**Movie S5 (separate file).**

Z slices of tubes reconstructed by HVEM

**Movie S6 (separate file).**

Rendering of tubes reconstructed by HVEM

**Movie S7 (separate file).**

FISH observation of the tubes without nuclease treatment. Z range; 2.78 μm, Z step; 0.31 μm.

**Movie S8 (separate file).**

FISH observation of the tubes after DNase treatment. Z range; 1.31 μm, Z step; 0.15 μm.

**Movie S9 (separate file).**

FISH observation of the tubes after RNase treatment. Z range; 1.28 μm, Z step; 0.15 μm.

